# Efficient Transmission in the Blowfly Early Visual Synapses Through the Regularization of Vesicle Release

**DOI:** 10.64898/2026.06.11.731739

**Authors:** Golam M Kashef, Rob de Ruyter van Steveninck

## Abstract

Early studies of synaptic transmission by Bernard Katz and colleagues suggested that neurotransmitter release at graded-potential synapses occurs through statistically independent (i.e. Poissonian) quanta [1, 2]. Subsequent experimental work supported this framework [3]. However, these measurements were performed in vitro on relatively simple synapses and under non-physiological conditions, often converting spiking neurons into graded-potential neurons through the use of channel blockers. Relying on the conventional assumption that vesicle exocytosis follows a Poisson process, measurements of the contrast power transfer spectrum and noise power spectral density of large monopolar cells (LMCs) in the blowfly *C. vicina* imply a sustained vesicle release rate exceeding 10^5^ vesicles per second per LMC. Given the physical dimensions of photoreceptors and synaptic vesicles, such a release rate appears physiologically implausible. If vesicle release is more temporally structured, low-frequency noise could be suppressed, substantially reducing the vesicle release rate required to account for experimental observations. The reduction of noise at low frequencies is especially advantageous given inputs such as photoreceptor signals which are already low-pass filtered. Visual activity generates substantial extracellular potentials within the lamina cartridge [4]. We propose that these extracellular potentials regulate vesicle release by modulating the voltage sensors that trigger exocytosis. We provide experimental evidence for the connection between currents driving the LMC and the extracellular potentials during visual activity, and demonstrate, using simple models, how effective “Poisson” rates are maximized due to vesicle regularization.

---

From transcription and translation in protein synthesis, through quorum sensing in bacteria, to neural control of behavior, and even to social interactions within and between multicellular species, biological systems depend on signaling across multiple scales to sustain themselves. In almost all such cases, signals must contend with degradation arising from thermal and other forms of noise. Mechanisms that improve signal transmission are diverse, but improvements in information transfer invariably come at a metabolic cost, directly or indirectly. Examples include making several copies of a genetic code, ramping up of signaling molecule production [5], and the formation of myelin sheaths around axons involved in long-distance communication-all of which are energetically demanding. Moreover, spatial constraints become relevant because resources that are not immediately used must be stored physically until they are expended. Because of these constraints, information transmission has to be *resource efficient* in order to be viable.

The signaling system we are exploring here is an array of chemical synapses connecting six photoreceptor axons to a pair of second order visual neurons called the Large Monopolar Cells (LMCs) in the early visual system of the blowfly *Calliphora vicina*. Signal transfer from photoreceptor to LMC is mediated by vesicles released from synapses located at the photoreceptor axon. Since synaptic vesicles are essentially quantal packets that carry relatively specific amounts of transmitter, we should think of signaling in chemical synapses as being mediated through discrete events. The properties of quantal transmitter release were first studied by Bernard Katz and colleagues[1, 2]. Spontaneous events (observed as tiny postsynaptic potentials of with a typical wave-form) were seen to arrive as a Poisson process with exponentially distributed interarrival times and normally distributed amplitudes with a coefficient of variability of around 25%[1, 2]. The distribution of amplitudes of the postsynaptic pulses as a response to short identical presynaptic pulses were seen to closely match this distribution convolved with a Poisson distribution, indicating that the number of events induced by a single pulse is Poisson distributed [1, 2]. These experiments were performed in the frog neuromuscular junction, pharmacologically treated to prevent spiking.

The results of Katz and colleagues create a picture where vesicles are released independent of each other in graded-potential synapses in a rate modulated Poisson process, with a typical (if noisy) postsynaptic response to a single release event. Mathematically, this is very similar to photon absorption in photoreceptors [6, 7] when exposed to modulated light intensity. One experimental method to estimate the rate of photon absorption at a blowfly photoreceptor involves exposing it to multiple repeated presentations of the same light contrast signal with a small standard deviation and flat spectrum (described further in the Methods section)[7–9]. By the analysis presented in de Ruyter and Bialek (2001) [9], we measure an “effective” Poisson rate (*R*_eff_) by dividing the contrast power transfer function by the average noise power spectral density and finding the maximum. The method is briefly summarized in Appendix A. Here we apply this logic to LMCs, replacing photon absorption events with vesicle release events.If we assume that vesicle release is described by a Poisson process, then a measurement of *R*_eff_ provides us with a lower bound on the release rate. Any additional noise sources in the pro-cess, such as variation in quantal content, photon noise etc., will only increase that lower bound.

In a lamina cartridge, the axons of six photoreceptors (R1-R6) from different ommatidia (but all collecting light from the same direction) converge onto a pair of LMCs (L1-L2) and each of the roughly 1320 active zones in the photoreceptor axons (220 per photoreceptor) simultaneously synapse onto both the LMCs through specialized “tetrad” synapses, leading to parallel processing of photoreceptor signals by the LMC pair [4, 10–13]. Weckström and Laughlin (2010) studied potentials generated in the extracellular space during visual activity and suggested that these play a role in shaping the LMC response[4] (see Figure 1). This is important because synaptic vesicle release is mediated by voltage gated Ca^2+^ channels in the presynaptic active zone, whose rate of opening depends on the photoreceptor *transmembrane* voltage.

**FIG. 1.**
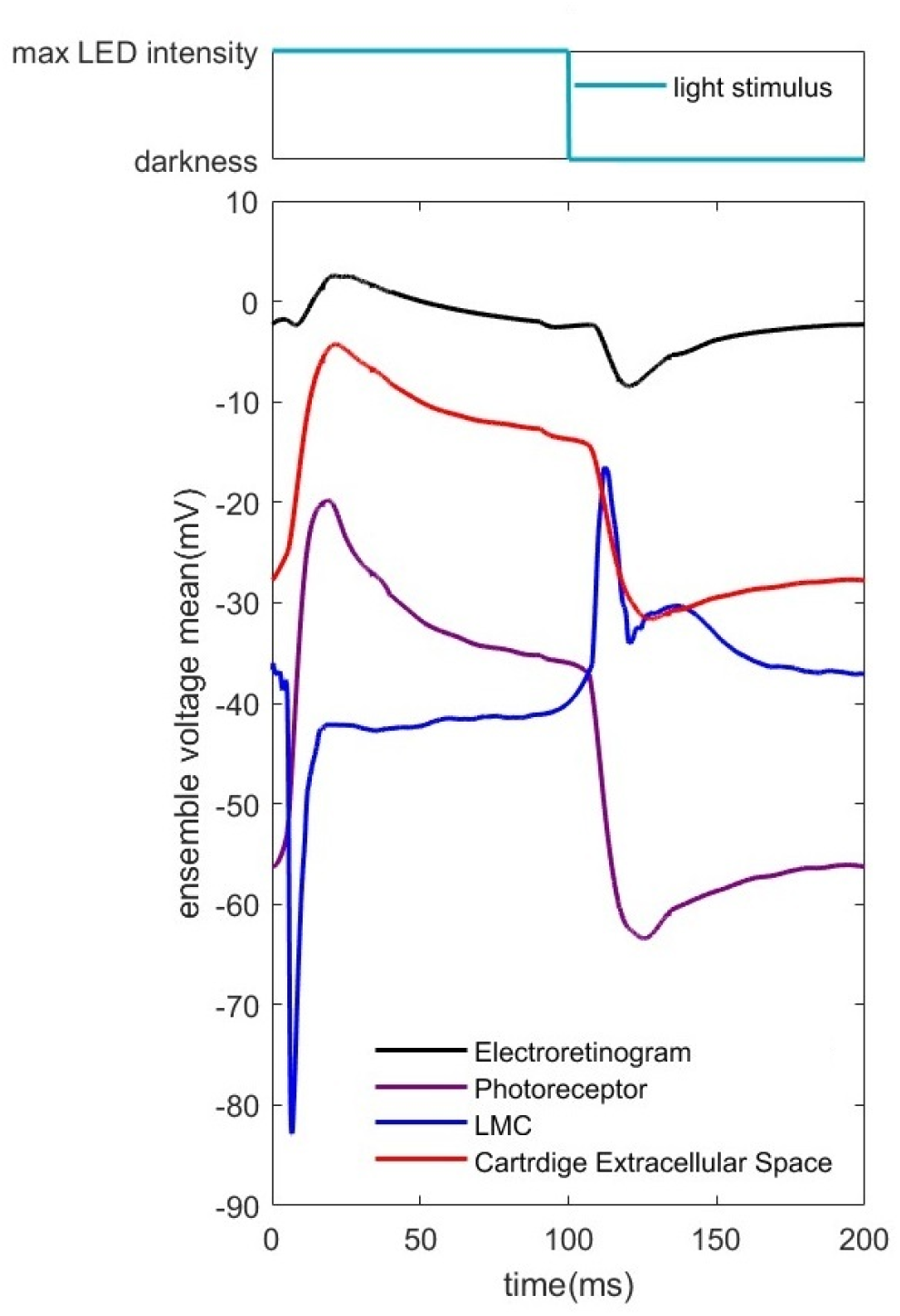
**Top:** The light intensity of the LEDs as a function of time during a repetition of the pulsed light stimulus. The pulsed light is actually a 5Hz square wave, with the LED at maximum brightness for 100ms, followed by 100ms of darkness. **Bottom:** Measured background electroretinogram, along with photoreceptor axon, LMC and cartridge extracellular potential measured in the same cartridge. The amplitudes and waveforms of the signals agree with what we know from literature.

Here we study the signals generated in the extracellular space in parallel with those in the photoreceptor and LMCs. We perform an experiment where carefully designed random currents are injected through the electrode while a light contrast signal is being presented, in order to compute the current-to-voltage impulse response functions. We propose that this leads to regularization of vesicle release, and show using simple models how this might explain high effective Poisson rates observed in LMC responses.

## I. MATERIALS AND METHODS

### A. Collection and preparation of flies

The *C. vicina* used in our experiments were collected from Dunn Woods at Indiana University during spring-time. In order to prepare a fly for an experiment, it is placed in a tube with its head exposed, and affixed with wax. A small hole is cut in the right eye for entry of the recording electrode tip, near the top and at a position that allows for easy targeting of the lamina. This is sealed with silica gel to prevent the hole from drying out, and to minimize parasitic capacitance. An incision is made near the bottom of the contralateral eye for the entry of the reference electrode. The tube containing the fly is attached to a post mounted on a pair of goniometric stages, one to control the pitch angle, and one for yaw.

### B. Electrodes preparation, properties and usage

Recordings are made using the Axoprobe-1A microelectrode amplifier. Borosilicate glass micropipettes (1.2mmOD 0.6mmID) are fabricated using the Sutter Model P-87 micropipette puller, using slightly lower heat settings than recommended for intracellular recordings to produce lower resistance electrodes, since lower resistance electrodes are better for current injection experiments. The resistances of the electrodes are between 80-100MΩ. The fast nature of our current injection signals interferes with the capabilities of the bridge balance function of the amplifier, so we do not use the bridge balance. Instead, after a series of current injection experiments, we very gradually retract the electrode until the tip is at a shallow part of the track, and perform the same experiment at “ground”. The impedance to ground is calculated by subtracting the impedance at ground from the raw calculated impedances. There is an effective parallel capacitance associated with current injections whose effects are noticeable at high frequencies- this capacitance is removed numerically from the electrode and all other measured impedances before this subtraction (the method was tested and works well in electronic components with impedances mimicking membrane R|| C values from literature).

For our electrode solution we use 2M K-acetate and 4mM KCl, following Weckström and Laughlin (2010)[4].

### C. Finding cells

A track is made by advancing the electrode tip vertically using a manual mechanical microcontroller designed for very fine adjustments. The goniometric stages are used to adjust the pitch and yaw angles of the fly in order to get the best track for finding lamina cartridges. Once the electrode is inside the fly, the voltage is set to zero, and a moveable array of blue-green LEDs (whose wave-length matches the peak spectral sensitivity of blowfly photoreceptors) pulses light as a 5Hz square wave. The signals are monitored on an oscilloscope. The electrode is advanced gradually with strategic adjustments to LED placement, until the waveform associated with the LMC is seen on the scope (Figure 1, blue trace). The position of the LEDs is further adjusted until the central LED is at the center of the cell’s point spread function, where the amplitude of the waveform is maximized.

When in a lamina cartridge, creating a light mechanical disturbance or an extremely fine adjustment of the microcontroller can cause the electrode to penetrate a photoreceptor or enter the extracellular space in the same cartridge. The light induced signals in combination with the resting potentials in these recordings are characteristic of each recording location (see Figure 1). Using another fine adjustment, we can move from the extracellular space to a photoreceptor or vice versa.

### D. Delivering the light contrast signal and finding effective Poisson rates

Both the light contrast and current injection waveforms are 500ms long (the frequency resolution is thus *δf* = 2Hz) and delivered through a digital to analog converter to an LED driver circuit at a sampling rate *f*_sample_ = 5kHz, using data from MATLAB scripts. The light contrast signal is generated by adding sine waves of the same amplitude for all frequencies between *δf* and some maximum frequency separated by *δf* . Each sine wave has a random phase shift taken from a uniform distribution between *− π* and *π*. The mean is subtracted, and the signal is multiplied by *σ*_*c*_*/σ* where *σ*_*c*_ = 0.25 is the contrast standard deviation we want to set, and *σ* is the calculated standard deviation of the sum of the sine waves. This final product is a contrast signal with perfectly flat spectrum up to a maximum frequency. This is stored as a MATLAB data file, and used for all experiments in this work.

For measuring the effective Poisson rate, 200 repeats of the LED signal are presented to the fly, and voltages recorded are stored as MATLAB data files. The ensemble average of the 200 voltage traces is thus the *signal*, and deviations of the individual traces from this signal represent the *noise*. The ratio of the contrast power transfer function and the noise power spectral density gives us an effective Poisson rate, whose peak gives us the minimum rate of quantal arrivals needed to explain the observed signal and noise given the contrast. This is explained in Appendix A (see also de Ruyter and Bialek, 2001[9]).

### E. Experiment with simultaneous light and injected current stimulation

In order to understand the electrical properties of the system as it is being shown a light contrast signal, we study the responses of the photoreceptor and LMC intracellular space, and the cartridge extracellular space to current injections while it is simultaneously subject to the light contrast. While experiments involving current injections are commonplace, what is different about our experiment is that instead of pulsing the current as a boxcar function, we inject the current as a carefully prepared preconceived series of 200 different random signals (of the same length as our light contrast signal), symmetrically distributed around *I*_inj_ = 0 with standard deviation *σ*_*I*_ = 0.05nA.

Figure 2 shows the basic schematic of our current injection experiment. Here, the stored light contrast from Section D is simultaneously presented with injected current. Since we have to make the comparisons in Figure 6, we perform this experiment in the same cell/environment immediately after performing the previous one. We prepare *N* = 200 injected current signals *I*_*i*_(*t*) of period 500ms, with symmetrically distributed values around mean *µ*_*I*_ = 0 and standard deviation *σ*_*I*_ = 0.05nA. We calculate the current-to-voltage impulse response function *z*(*τ*) as the temporal cross-correlation between *V*_*i*,current_(*t*) (see Equation 2) and *I*_*i*_(*t*), multiplied by 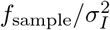, i.e.

**FIG. 2.**
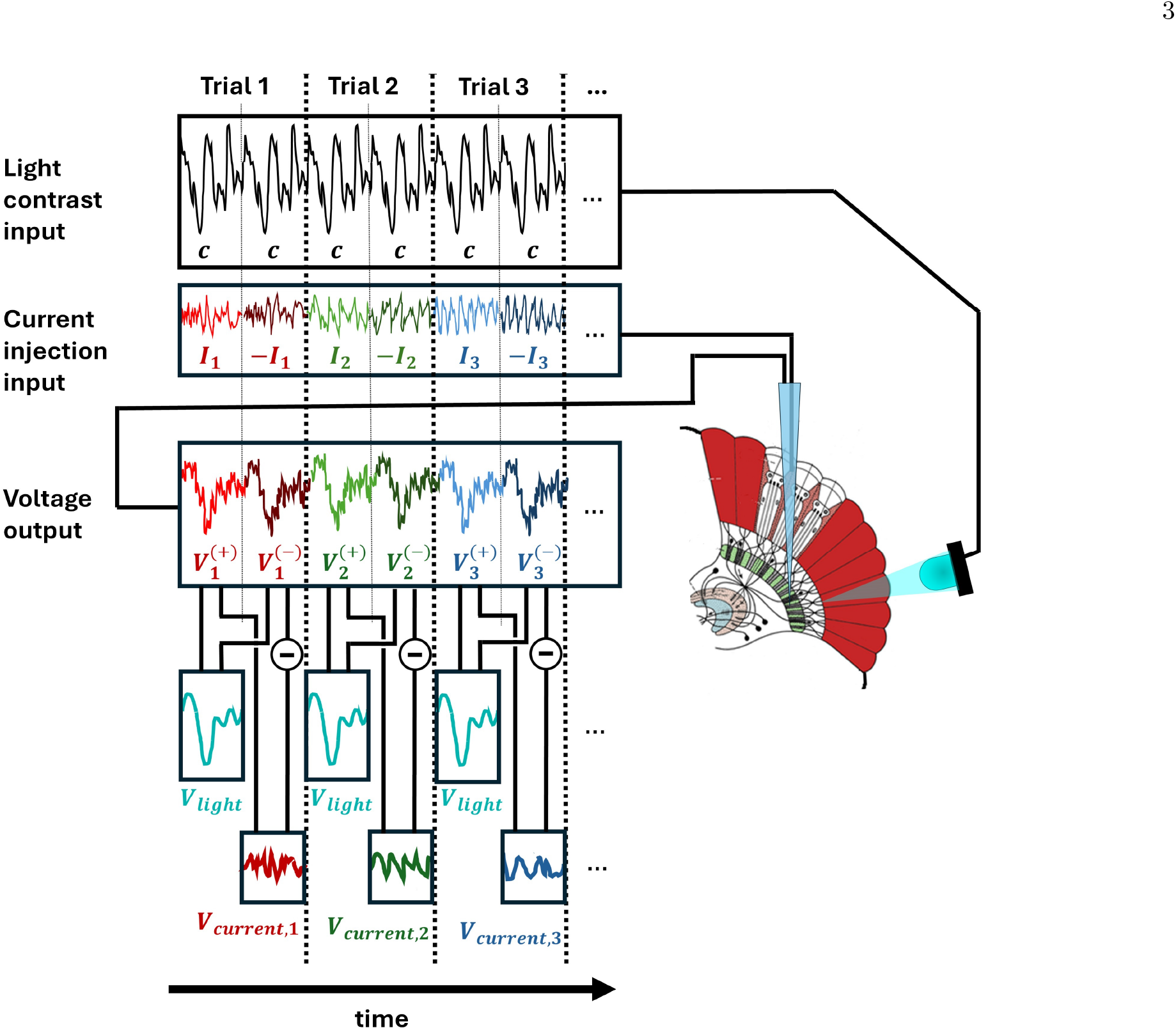
Setup of experiment with simultaneous light stimulation and current injection inside different points in the blowfly lamina cartridge. The same light contrast is used over all trials, and two repeats of it are presented within a each trial. A different injected current trace is used in each trial. Two repeats of the same trace are presented inside a trial, but with the sign of the second repeat inverted. If responses to both are linear, the resulting voltage output now has a part induced by the light contrast, and one induced by current. The former can be found by adding half the response to the sign inverted current to half the response to the original current. The latter can be found by subtracting half the response to the sign inverted current from half the response to the original current.

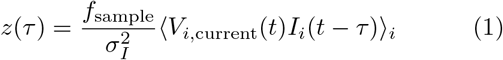

This assumes that the current signals are truly temporally uncorrelated. For a limited number of presentations *N*, this is strictly true when

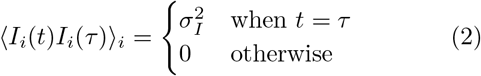

or, in other words,

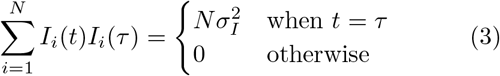

for all *t*. If we prepare our set of injected current signals as a matrix of *N* rows where the *i*^th^ row represents *I*_*i*_(*t*) (the *n*^th^ column represents the signal at time *t* = *n · δt*),then Equation 3 demands *orthogonality* between all the columns of the matrix. This is, of course, impossible if the number of columns of the matrix is greater than *N*, which represents *N/f*_sample_ = 40ms. However, since *z*(*τ*) dies out long before *τ* = 40ms (see Figure 7), nonorthogonality between any pair of columns separated by *N −*1 or more columns does not affect our analysis. More-over, if we repeat the same order of *N* = 200 orthogonal columns over and over until the number of columns represent the required period of 500ms (i.e. 2500 columns), every pair of columns separated by any number of columns other than *N −* 1 times an integer is an orthogonal pair.

With this knowledge, we first generate a pseudorandom 200 *×*200 matrix, where each element is drawn from a Gaussian distribution with mean 0 and standard deviation 1. We then computationally perform singular value decomposition on this matrix [14], giving us another 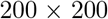 matrix with mean 0 and standard de-viation 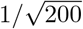, this time with orthonormal columns. The distribution of the entries is symmetric around 0, but not exactly Gaussian. Multiplying this matrix by *σ*_*I*_ 200 gives us a matrix with orthogonal columns, and where the entries have mean 0 and standard deviation *σ*_*I*_ = 0.05nA. The rows of this matrix are each of the *I*_*i*_(*t*) we use for current injection. The signal is scaled appropriately and sent to the Axoprobe amplifier which converts it to current injected through the electrode.

As represented in Figure 2, each *I*_*i*_(*t*) is presented twice, once in its original sign and once with the sign inverted. Simultaneously, our single stored light contrast signal is presented with all presentations of current signals, keeping the same polarity each time. Let us define each of the voltage traces when the *i*^th^ injected current signal is presented in the original sign as 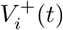 and each of the voltage traces when the *i*^th^ injected current signal is presented with sign inversion as 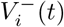. This allows us to explicitly decompose our voltage traces into the signal that comes from light stimulation and the signal that comes from current injection, as follows:

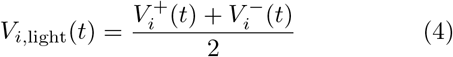

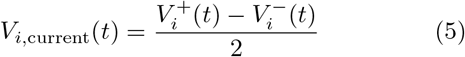

Equations 4 and 5 assume that the voltage response generated by current in one direction is equal and opposite to the voltage response generated by the current in the opposite direction, a hallmark of linearity. The way to check that this is indeed the case is to compare the ensemble average of *V*_i,light_(*t*) to ⟨*V* (*t*)⟩, the ensemble averaged voltage signal *without* current injection.

## II. RESULTS

### A. Responses of Photoreceptor, LMC and Cartridge Extracellular Potentials to Pulsed Light

Figure 1 shows the photoreceptor, LMC and cartridge extracellular potentials in response to typical pulsed light (set to the maximum brightness allowed in our setup) measured in our setup from the same lamina cartridge (ensemble averaged over 50 repeated presentations). Also shown is a typical background electroretinogram, a small signal with a wide point spread function found when the tip of the recording electrode is embedded in the retina extracellular space (note that it is more or less close to the “ground” voltage).

These waveforms agree with signals we know from literature[4, 7]. The photoreceptor follows the light intensity with a slight delay, an overshoot at the ON signal and a somewhat smaller undershoot at the OFF signal. The LMC signal is sign inverted and adapts quickly to the background light intensity (acting like a differentiator), with a strong ON and a somewhat less strong OFF signal. The cartridge extracellular potential (explored by Weckström and Laughlin [4]) is a smaller and slightly delayed version of the photoreceptor signal. The resting voltages of the photoreceptors, LMCs and the cartridge extracellular potentials are found to be *−* 55mV*−* 60mV, *−* 35mV*−* 40mV and *−* 25mV*−* 32mV respectively, also matching what we know from literature [4, 15, 16].

### B. Contrast Impulse Response Functions of Photoreceptor, LMC and Cartridge Extracellular Potentials and Measured Effective Poisson Rates

Potentials are measured from a photoreceptor, LMC and cartridge extracellular space from the same lamina cartridge for 200 different repetitions each of the same *σ*_*c*_ = 0.25 flat spectrum contrast, with mean intensity equal to half of the maximum possible LED intensity. Figure 3 shows the linear contrast impulse responses of the ensemble average of these three potentials in units of millivolts per unit contrast per millisecond. The photoreceptor contrast impulse response is largely monophasic, with a peak at 10ms. The LMC response function is biphasic and akin to a delayed sign-inverting differentiator. There is a strong minimum (much larger than the photoreceptor peak) at around 10ms, and a maximum around 20ms. The extracellular impulse response function is a smaller, delayed version of the photoreceptor impulse response function.

**FIG. 3.**
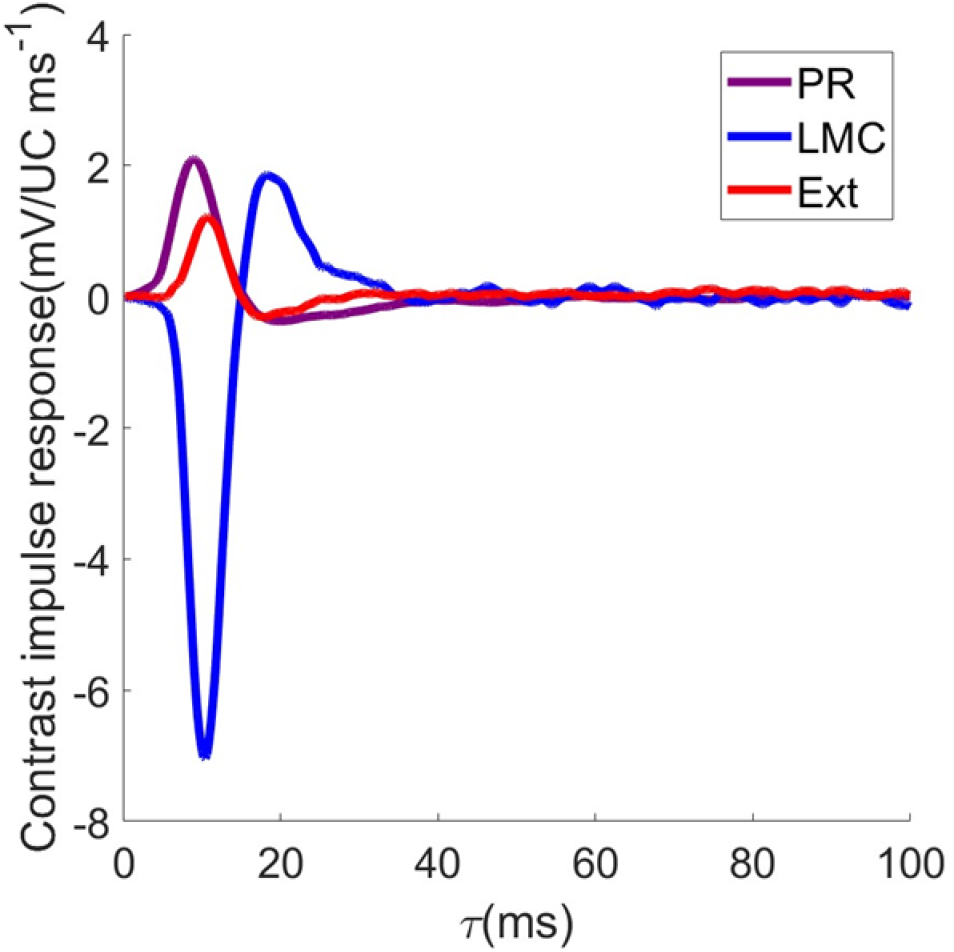
The contrast impulse response, computed from the measurements as 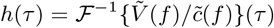 in units of milliVolts per unit contrast per millisecond for a photoreceptor, LMC and extracellular potential from the same lamina cartridge in response to a flat spectrum contrast. These waveforms represent the response of a cell to a brief 1ms pulse of light at *τ* = 0, with an amplitude equal to the mean light intensity in the experiment.

Figure 4 shows the contrast power transfer, noise power spectral density and the effective Poisson rate for the photoreceptor, LMC and cartridge extracellular potentials. For the photoreceptor, the maximum of *R*_eff_ is at approximately 2*×* 10^5^/s, meaning that this must be the minimum photon absorption rate that explains the signal and noise levels given the contrast function. For the LMC, this maximum is found to be around 5 *×*10^5^/s. The fact that this is higher than the photoreceptor effective Poisson rate is consistent with the fact that the LMC is downstream of the photoreceptor, since *six* photoreceptors representing the same direction in space are signaling to the LMC in a parallel fashion, meaning that perfect transmission would require the rate to be 12*×* 10^5^/s, which is higher than what we find here. If we are to assume that the LMC is driven by a rate-modulated Poisson process with independent vesicle release events, this 5 *×*10^5^/s value would be the minimum possible rate of vesicle release required to explain the data given Poissonian delivery of vesicles. With 1320 active zones, this is about 370 exocytosis events per second per active zone.

**FIG. 4.**
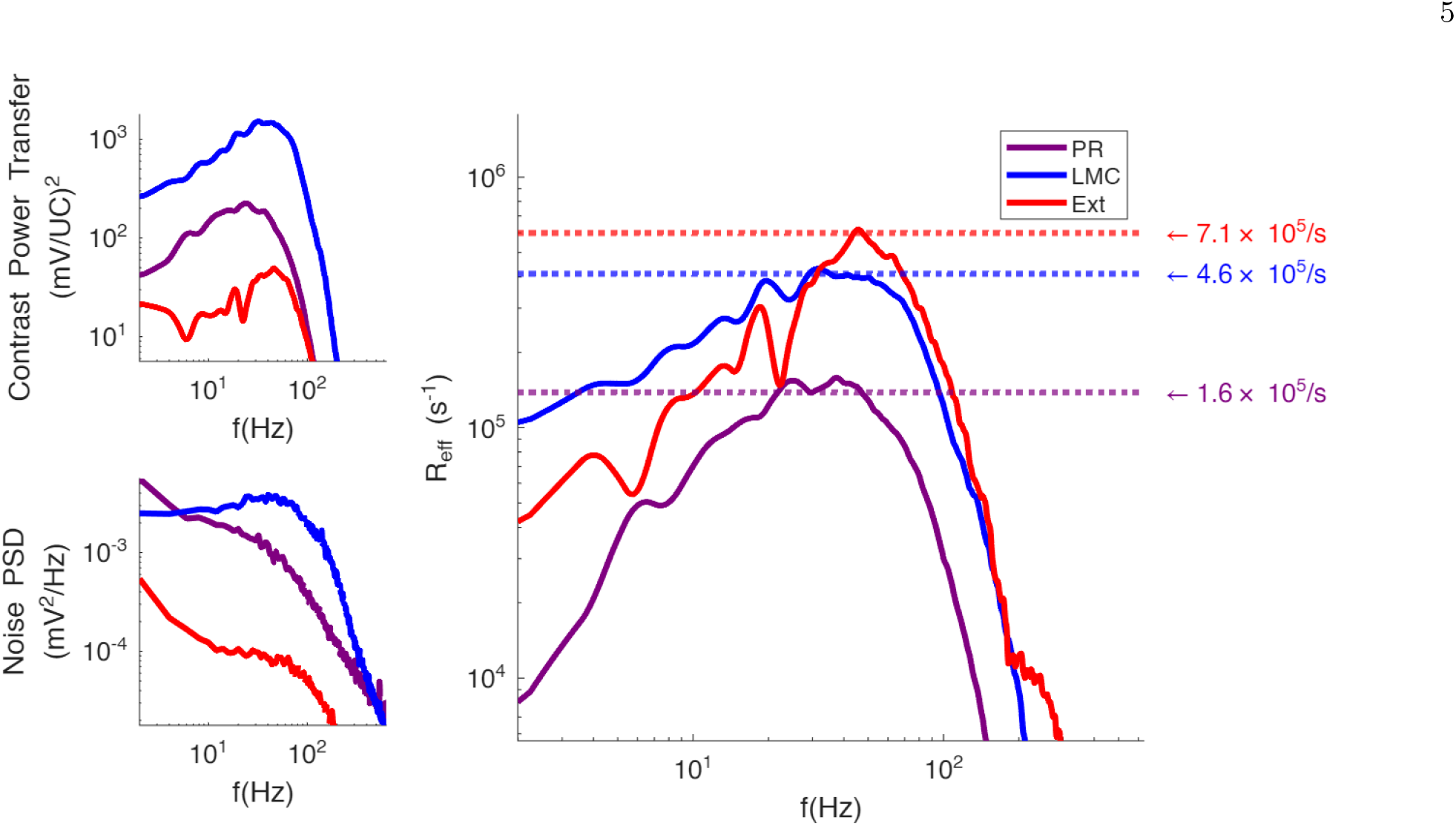
The contrast impulse response functions (A), noise power spectral densities (B) and *R*_eff_ (C) plots for photoreceptor, LMC and extracellular potentials from the same cartridge.*R*_eff_ is measured as 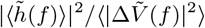 where 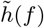 and 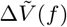 are respectively the Fourier transforms of the contrast impulse response and the deviations of individual voltage traces from the ensemble average. We find that the maximum of the effective Poisson rate is logistically implausible, given what we know about the geometry of photoreceptors and synaptic vesicles.

Interestingly, similar high effective Poisson rates are also observed in the cartridge extracellular potential, and notably, the observed values across cartridges are relatively close to LMC effective rates. This suggests the possibility that these potentials are connected, and that the same discrete process driving the LMC may be involved in signal generation in the cartridge as well. It also suggests that the extracellular space is approximately isopotential, because for these to work, all active zones should contribute measurably to the extracellular potential, thus pooling the effective Poisson rates from all six photoreceptors. Figure 5 shows the *negative* cross correlation between the measured LMC and cartridge extracellular potentials in a single cartridge (the sign inversion is introduced because of the sign inverting property of the LMC). The peak is effectively at *τ* = 0, showing that these potentials track each other quite closely.

**FIG. 5.**
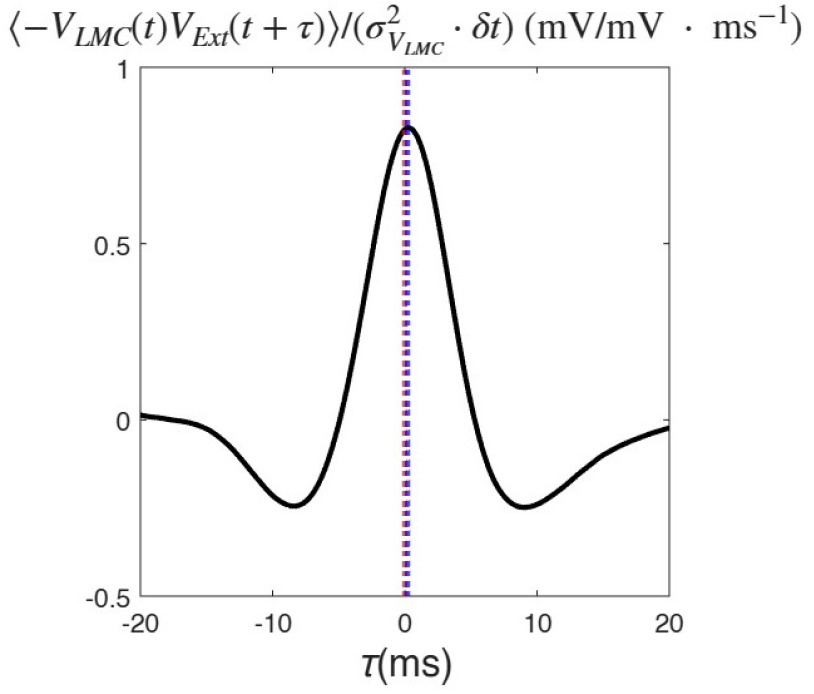
The negative cross correlation between the LMC and cartridge extracellular potentials. The time of the peak (represented with a blue dotted line) is 0.2ms, which is very close to *τ* = 0 (red dotted line). Note that 0.2ms is the sampling period of the data.

### C. Current Injection Experiments

As shown in Figure 6, the ensemble average of *V*_i,light_(*t*) matches *⟨V* (*t*)*⟩*, the ensemble averaged voltage signal without current injection with a tight fit around the line of equality, suggesting that we are in the linear regime. This is the case in our current injection experiment for the photoreceptor, LMC and cartridge extracellular potential, with an almost perfect match between the derived *⟨V*_i,light_(*t*)*⟩*_*i*_ and the signal with no injected current.

**FIG. 6.**
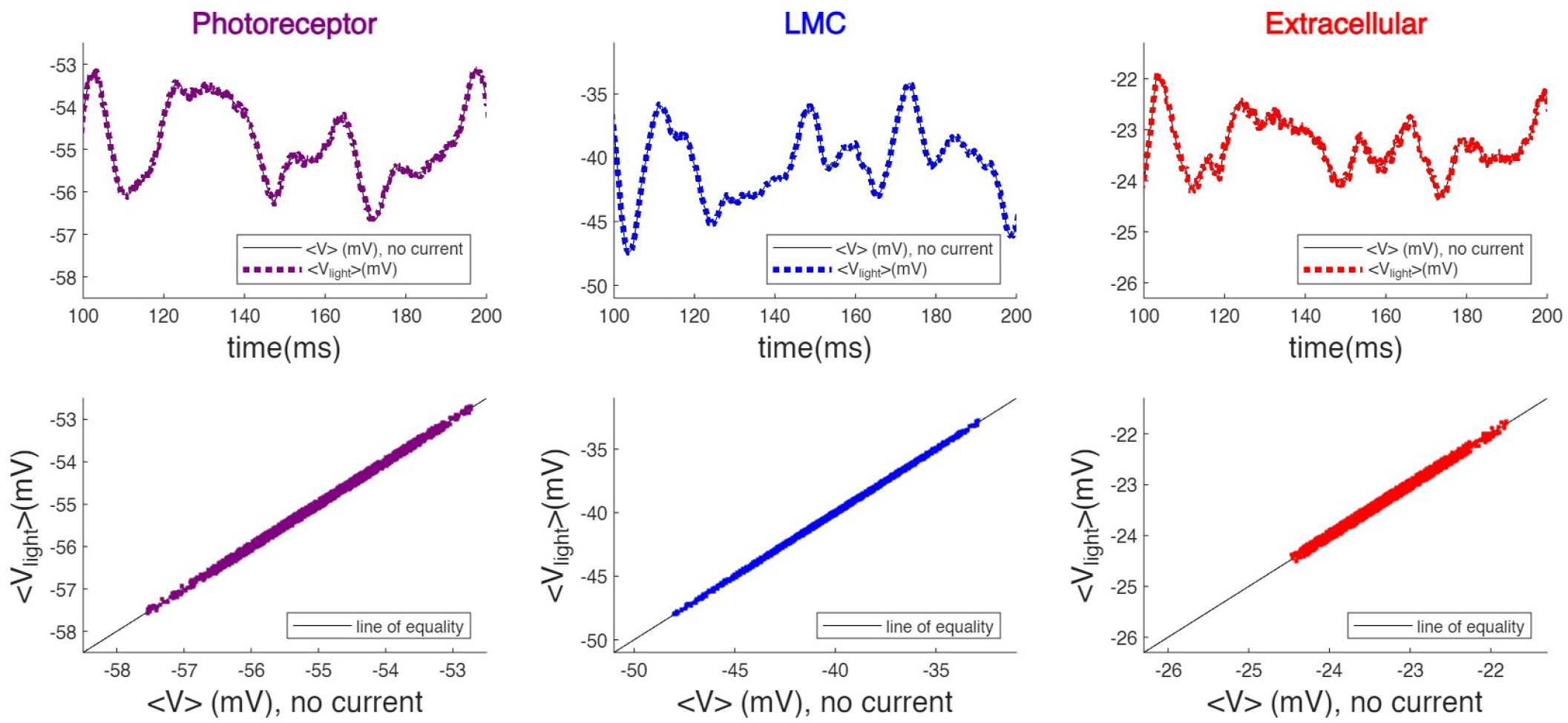
**Top panels:** Light induced signals in photoreceptors, LMCs and the cartridge extracellular space. The voltage signal in an experiment with no current injection is shown with a thin black line. Overlaid with dotted lines are the signals we get from the mean of the two signals with injected currents in opposite directions (see Equation 1). **Bottom panels:** Light induced voltage in experiments with injected current versus the counterparts with no injected current, plotted over lines of equality.

The current-to-voltage impulse response functions for photoreceptors, LMCs and the cartridge extracellular potentials are shown in Figure 7 in units of milliVolts per nanoAmps per millisecond. The complex impedances are simply the Fourier transforms of the current-to-voltage impulse response functions. The magnitudes of these are shown in the rightmost panel. As a first approximation, the best fit RC-filters are presented with dotted lines. The parameter values measured in different cartridges are in agreement with literature or are reasonably close (see Tables I and II)[4, 15, 17].

**FIG. 7.**
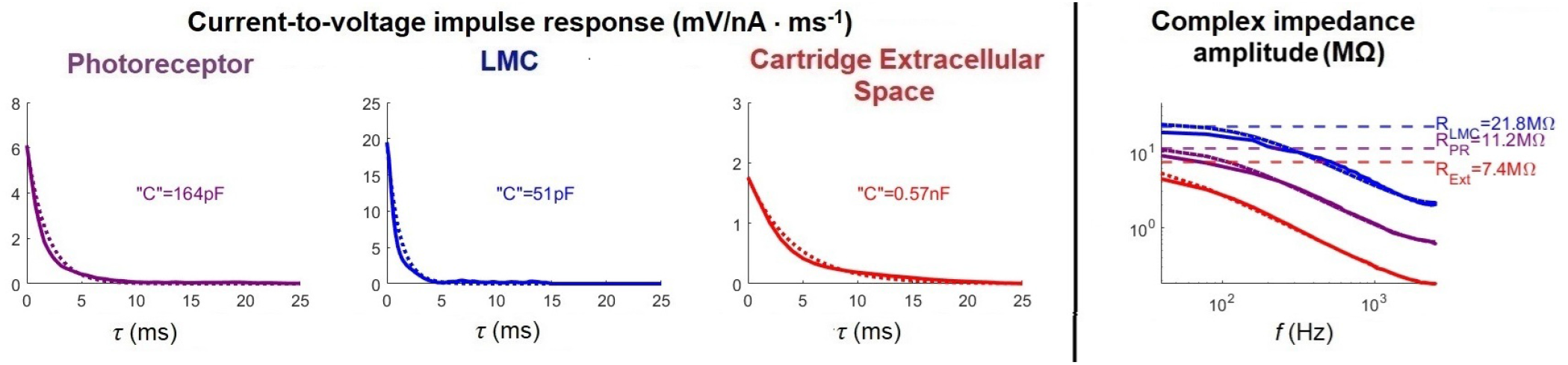
**Left panels:** The current-to-voltage impulse responses in mV/nA per ms (or MΩ per ms) for the photoreceptor, LMC and the cartridge extracellular space, with capacitance values (found from the reciprocal of the y-intercept) and best RC fit (dotted lines) shown. **Right panel:** Complex impedance amplitude in MΩ for the photoreceptor (purple), LMC (blue) and the cartridge extracellular space (red), with resistance values (found from the y-intercept) and best RC fit (dashed lines).

**TABLE I.** Resistance values of photoreceptors (N=2), LMCs and cartridge exatracellular spaces (N=3) found in our experiments from R || C curve fitting, compared with values from literature (found in light-adapted conditions unless stated otherwise)

| | Best-fit resistance from experiments (MΩ) | Resistance from literature (MΩ) |
| --- | --- | --- |
| Photoreceptor | $11 \pm 0.15$ | $8.75 \pm 3.2$ (Light-adapted),<br>$32 \pm 3$ (Dark-adapted)[17] |
| LMC | $21.8 \pm 1.3$ | $17.1$[15],<br>$20 \pm 4.4$ [18] |
| Cartridge extracellular space | $8.5 \pm 1.47$ | $7.8 \pm 0.78$ (Light-adapted),<br>$6.0 \pm 0.45$ (Dark-adapted) [4] |

**TABLE II.** Capacitance values of photoreceptors (*N* = 2), LMCs and cartridge extracellular spaces (*N* = 3) found in our experiments from R|| C curve fitting, compared with values from literature

|  | Best-fit capacitance from experiments (pF) | Capacitance from literature (pF) |
| --- | --- | --- |
| Photoreceptor | $167 \pm 3$ | 130 (Dark-adapted)[17] |
| LMC | $52 \pm 11$ | 43<sup>a</sup> |
| Cartridge extracellular space | $543 \pm 112$ | 510 (Light-adapted),<br>710 (Dark-adapted) [4] |

<sup>a</sup> This derived by multiplying the well known standard membrane capacity $10^{-5} \mu\text{F}/\mu\text{m}^2$ for neurons[19] with the total membrane area of LMCs found from van Hateren and Laughlin (1989)[15]: $900 \mu\text{m}^2$ for the synaptic zone, $1000 \mu\text{m}^2$ and $4333 \mu\text{m}^2$ for the axon (using axon length of $425 \mu\text{m}$ , and radius of $1.35 \mu\text{m}$ )

### D. Contrast to Current Impulse Response Functions

Now that we have impulse response functions for both contrast to voltage *and* injected current to voltage, we can use them to find a contrast to current impulse re-sponse function as

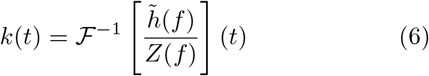

Where 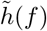 is the Fourier transform of the contrast impulse response function and *Z*(*f*) is the complex impedance. This is the theoretical impulse response function we would calculate if we had an ammeter to measure the conventional current injected into the recording site as a result of the light contrast stimulus. Figure 8 shows the photoreceptor, LMC and extracellular contrast to current impulse response functions from the same cartridge.

**FIG. 8.**
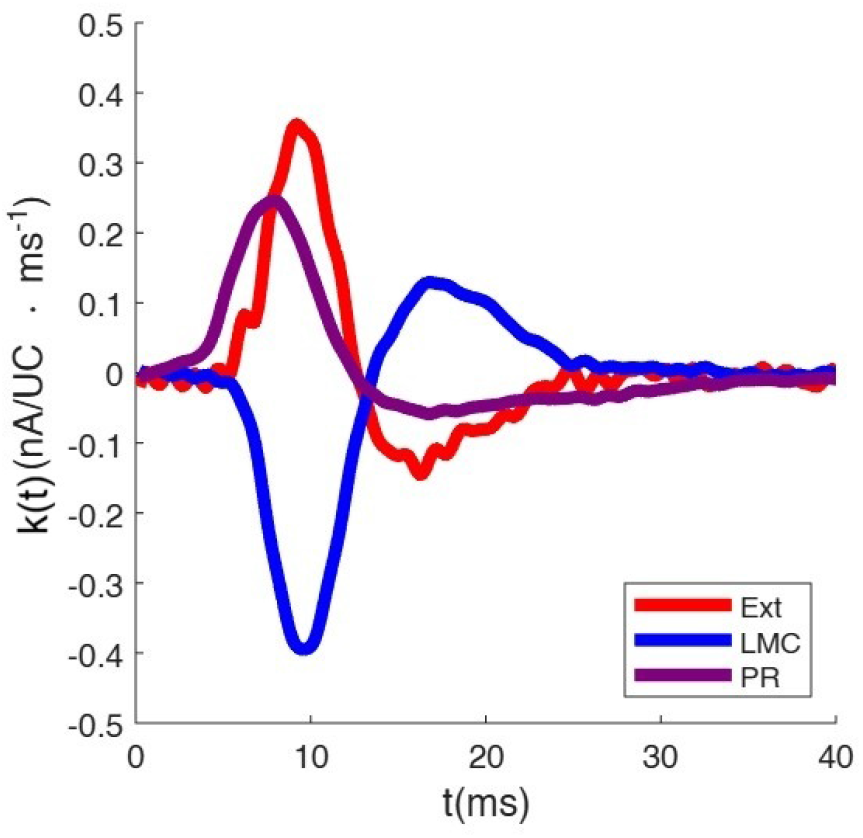
Contrast-to-current impulse response functions for extracellular, LMC and photoreceptor signals measured from the contrast impulse response function and the complex impedance.

What is perhaps interesting is that the extracellular contrast to current impulse response seems to almost match the LMC’s, but in the opposite direction. Figure 9 shows *− k*_Ext_(*t*) overlaid with*− k*_LMC_(*t*) for data from three different cartridges (cartridge 3 is the one we have been seeing so far in other figures). It also shows *I*_Ext_(*t*) overlaid with *I*_LMC_(*t*)- the theoretical currents that we get from convolving our contrast to current impulse response functions with the contrast. There is a striking match between the extracellular current responses and the negative LMC current responses to light contrast.

**FIG. 9.**
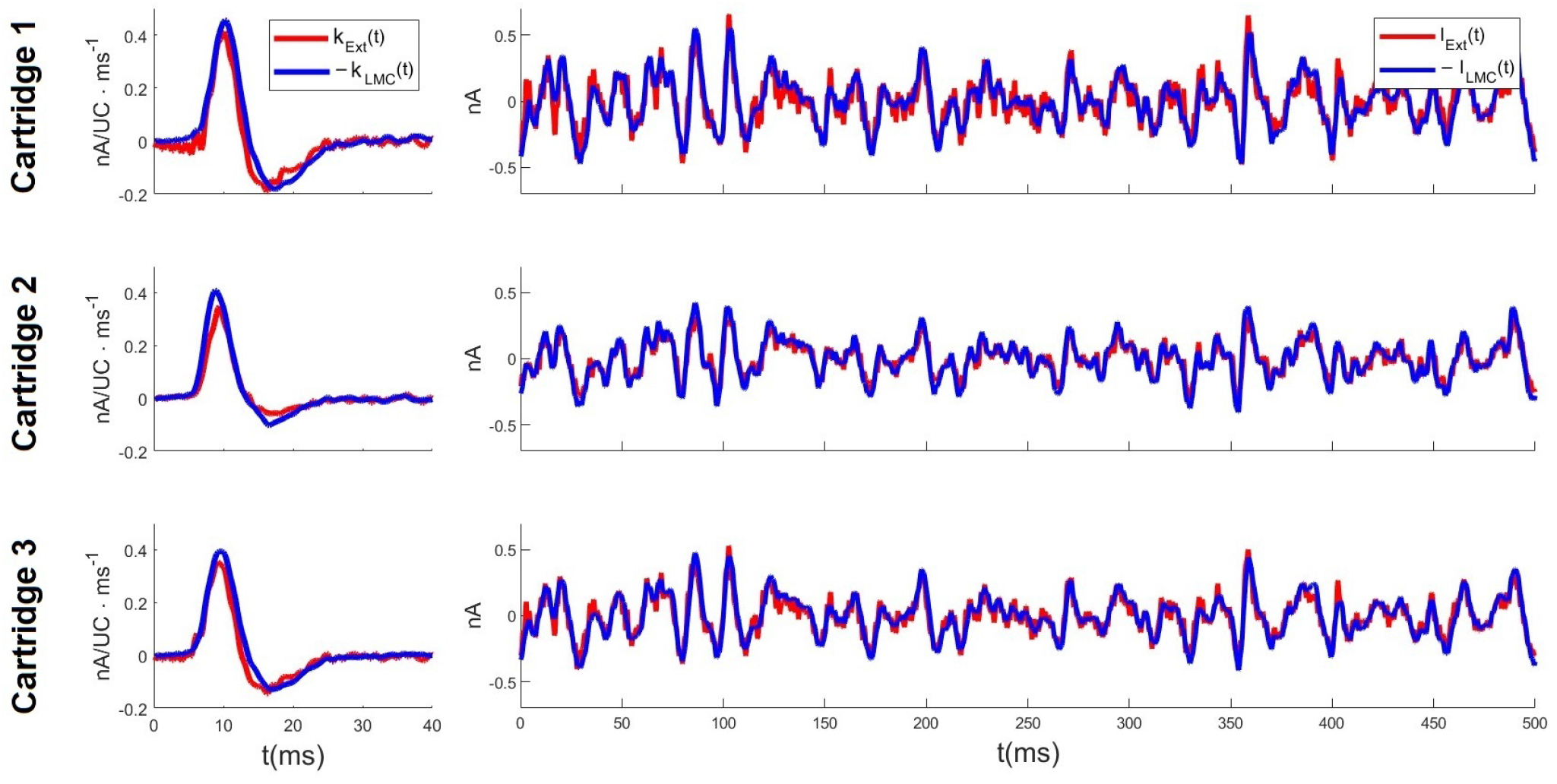
**Left panels:** Contrast-to-current impulse response functions for the extracellular space (*k*_Ext_) in three different cartridges, overlaid with the negative of its LMC counterpart (*k*_LMC_). **Right panels:** Theoretical current into the extracellular space (*I*_Ext_ = *k*_Ext_(*t*) *\* c*(*t*)) in three different cartridges, overlaid with the negative of the theoretical current into the LMC (*I*_LMC_ = *k*_LMC_(*t*) *\* c*(*t*)) during our light contrast experiment.

## III. DISCUSSION

The measured maximum effective Poisson rates for LMCs (Figure 4) agree with the rates found in previous studies performed under similar conditions [7], indicating a vesicle exocytosis rate of at least around 5*×* 10^5^ per second if the release events are statistically independent.

How feasible is such a high rate of vesicle exocytosis? Thinking of photoreceptor axons as cylinders, and noting that the part of the axon with active zones lies within the lamina cartridge, we may take the length of this part of the axon to be approximately equal to the length of the cartridge. This amounts to about *𝓁*= 50-60*µ*m [20– 22], which, along with the radius of *r* = 1*µ*m [13] gives us a total membrane surface area of 6 *·* 2*πr𝓁≈* 2000*µ*m^2^ for all six photoreceptors combined in the synaptic zone. A typical synaptic vesicle in the blowfly photoreceptor has a diameter of about *d* = 50nm [23–25]. This means that a typical synaptic vesicle has a membrane surface area of *πd*^2^ *≈* 7.8 *×* 10^*−*3^*µ*m^2^. This indicates that to maintain a sustained vesicle release rate of about 5 *×* 10^5^ s^*−*1^ (or greater), a total membrane area exocytosis rate of 4000*µ*m^2^s^*−*1^ must be achieved. In terms of the total surface area of photoreceptor membrane in the synaptic zone, this amounts to exocytosis of the total membrane area more than twice each second, a rather implausible feat. The calculation assumes a full range of modulation in the vesicle rate given the light contrast signal. However, even if the range was limited to 50%, this approximated lower bound on the vesicle rate would fall only by a factor of 4 (see Appendix A), and we would still be in a physiologically implausible range.

Going back to the results in Figure 8, what is the source of injected current arising from light stimulus? For photoreceptors, these are likely generated through the movement of positively charged ions into the photoreceptor as a result of the opening of TRP and TRPL channels. The opening up of these channels leads to about a threefold drop in the input resistance of the photoreceptor (see Table 52 of Weckström, Hardie and Laughlin, 1991[17]), but these channels are in the retina and not the lamina- indicating that other potentials in the lamina may not be affected or only affected in a limited capacity by this signal. For the LMC, these currents are generated by the opening up of histamine gated chloride channels, and negatively charged Cl^*−*^ ions entering the cell as a result of synaptic vesicles (containing histamine) in photoreceptor active zones exocytoting into the synaptic cleft.[10, 12, 24] The fact that Cl^*−*^ is negatively charged explains why the sign of the impulse peak is negative for the LMC (and why LMCs are sign inverting). Like the contrast-to-voltage impulse response, this current impulse response is also biphasic. Specifically, the synaptic current associated with chloride should be given by

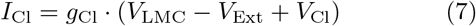

where *g*_Cl_ is the instantaneous chloride conductance produced by open chloride channels as a result of synaptic activity and *V*_Cl_ is the chloride associated reversal potential, which acts as a “battery” to draw chloride ions into the LMC. A typical intracellular chloride concentration is about 108 mM, whereas the extracellular concentration is approximately 560 mM [26, 27]. Substituting these values into the Nernst equation at 25°C (298.15K) yields the corresponding chloride reversal potential.

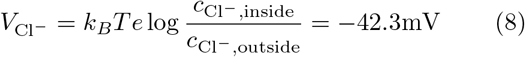

This is supported by voltage-clamp experiments reported by Laughlin and Osorio (1989). When a constant current is applied to hold the LMC at approximately 30–45 mV below its resting potential, the polarity of the LMC light response reverses.[18]. In Figure 10, we plot the current *I*_LMC_(*t*) from cartridge 3 (see Figure 9), shifted forward and backward in time by integer multiples of the sampling interval, against *V*_LMC_(*t*) *− V*_Ext_(*t*) + *V*_Cl_. We observe that for a time shift of *δt* = 0.8ms (corresponding to four sample periods), the two quantities strongly exhibit a linear relationship. The same pattern is consistently observed across other cartridges. The slope of the best linear fit, which has units of conductance, is 47.9 *±*4.3nS. The emergence of this relationship between the current and the LMC transmembrane potential on a submillisecond timescale supports the interpretation of *I*_LMC_ as the synaptic chloride current.

**FIG. 10.**
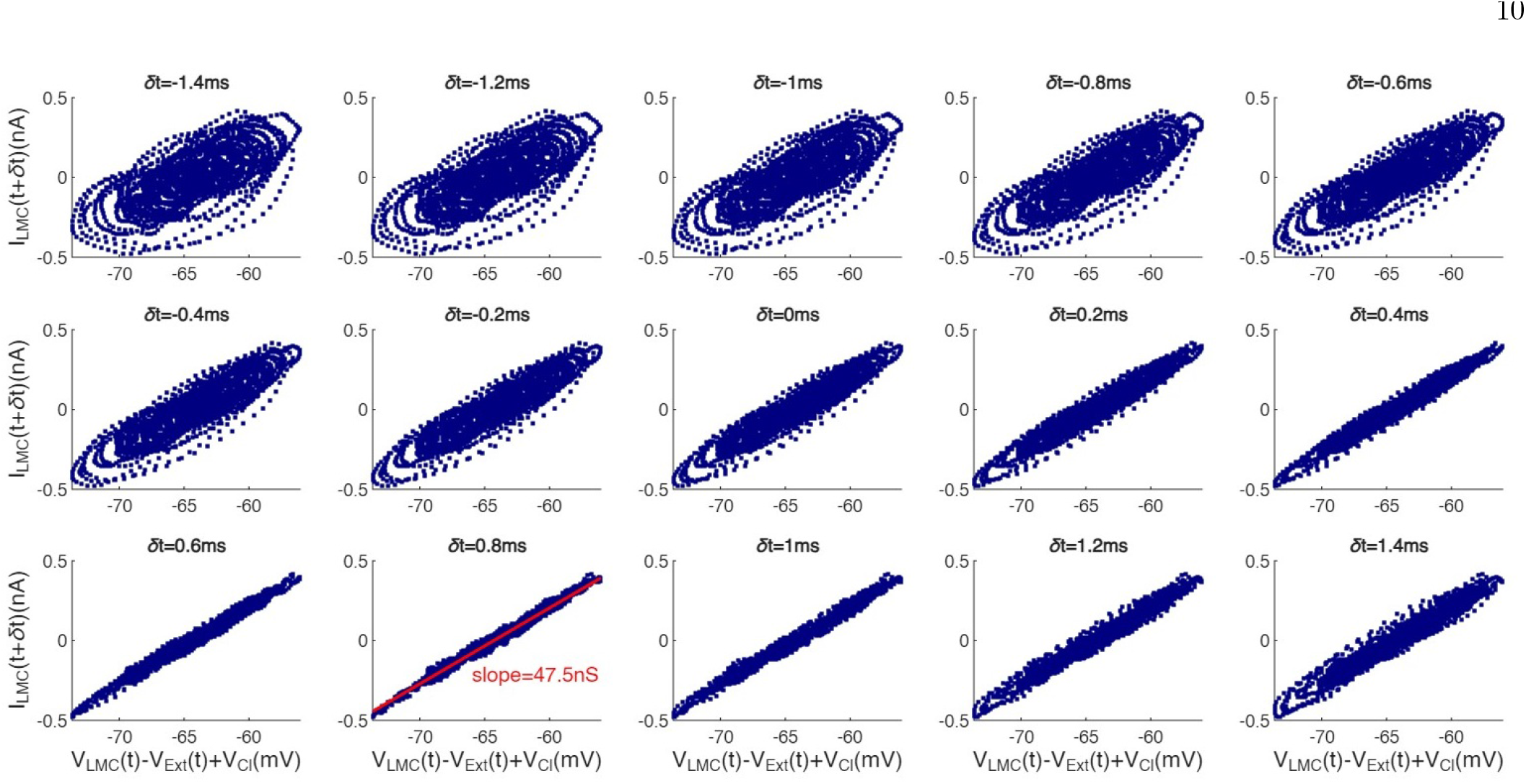
Theoretical *I*_LMC_(*t*) shifted forward and backward in time by integer multiples of the sampling interval, against *V*_LMC_(*t*) *− V*_Ext_(*t*) + *V*_Cl_ from the same cartridge. At a time shift of *δt* = 0.8ms (corresponding to four sampling periods), the two quantities display a tight linear relationship. The slope of the best linear fit is 47.9 *±* 4.3nS.

The match between the extracellular current responses and the negative LMC current responses to light contrast is perhaps not too surprising since the Cl^*−*^ ions that are taken up the LMCs are drawn from the cartridge extra-cellular space. What *is* rather surprising is that extracellular current responses match the negative LMC current responses and not *twice* it, since a *pair* of LMCs are drawing Cl^*−*^ ions during activity and not just one. The reason for this discrepancy is subject to speculation. A likely explanation for this is that the extracellular space is not purely a single compartment, which means that a portion of the distributed current into the space is shunted off, and does not contribute to the voltage at any one point.

However, if we are to take for granted that that the extracellular current to voltage impulse response is the result of synaptic chloride current, we could have a mechanism to induce a negative feedback in vesicle release (Figure 11). A single vesicle release causing a momentary opening of chloride channels and allowing chloride ions to be admitted into the LMC not only lowers the LMC voltage, but causes the extracellular voltage to go *up*. But since the rate of vesicle release events depends on the photoreceptor *transmembrane* voltage and this increase in extracellular voltage lowers this transmembrane volt-age, this causes the rate of vesicle exocytosis to fall temporarily. In preventing/slowing down the further release of vesicles, the extracellular potential relaxes, causing the transmembrane voltage and the release rate to go back up. The negative feedback caused by vesicle induced in-crease in the extracellular potential lowers the likelihood of an event immediately following another. In this framework, the occurrence of an event transiently suppresses the local rate, creating conditions under which successive vesicle exocytosis events are no longer statistically independent. Temporal correlations between events enhance predictability, as increased regularity in event timing allows the underlying rate to be inferred from fewer trials. This effect is particularly pronounced when observations are made over time windows longer than the typical inter-event interval. One can imagine here that in such a scenario, a relatively small (and more feasible) rate of vesicle release can lead to a high effective Poisson rate. This is explored in Kashef (2025)[28], which is also where this work is originally presented.

**FIG. 11.**
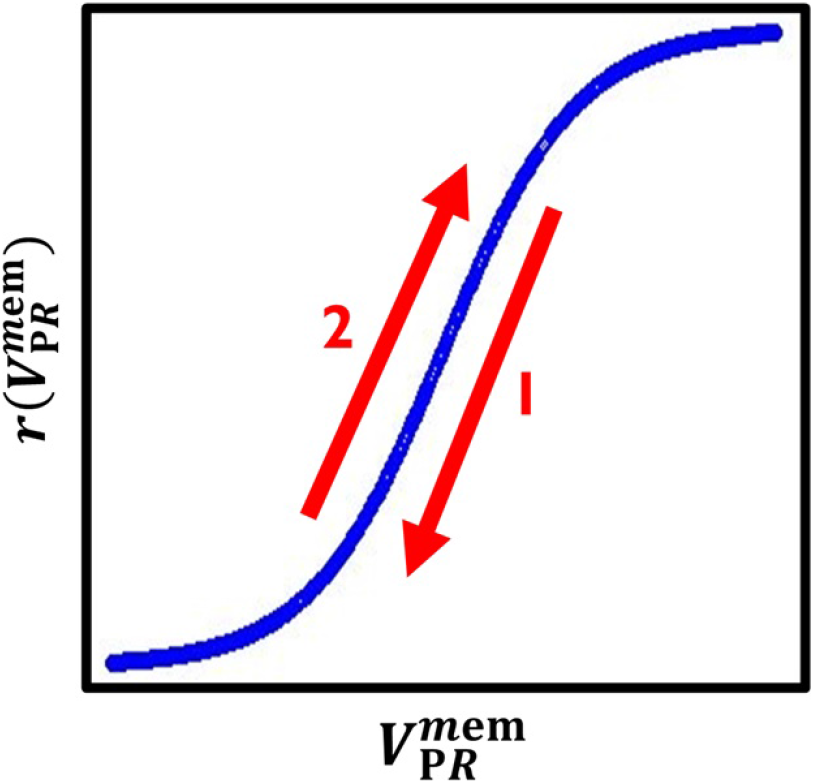
The rate of vesicle exocytosis *λ* is positively dependent on the transmembrane voltage 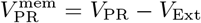. A possible negative feedback loop causing regularization of vesicles has the following steps (numbered on the plot). **(1)** High 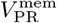causes transmitter release rate to go up, opening up chloride channels and raising *V*_Ext_, causing 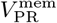 to drop. **(2)** The drop in 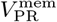 causes transmitter release rate to drop, lowering *V*_Ext_ and raising 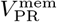 back up.

Because the relationship between the photoreceptor transmembrane voltage and vesicle release rate cannot be extracted directly from the data, we instead examine the relationship between the photoreceptor transmembrane voltage and the chloride conductance. This substitution is motivated by the expectation that the mean chloride conductance scales proportionally with the mean vesicle release rate. Using Equation 7 and assuming *I*_LMC_ = *I*_Cl_, we estimate the conductance *g*_Cl_ as *I*_LMC_*/*(*V*_LMC_ *− V*_Ext_ + *V*_Cl_). Figure 12 plots this quantity against the photoreceptor transmembrane voltage for cartridge 3.

**FIG. 12.**
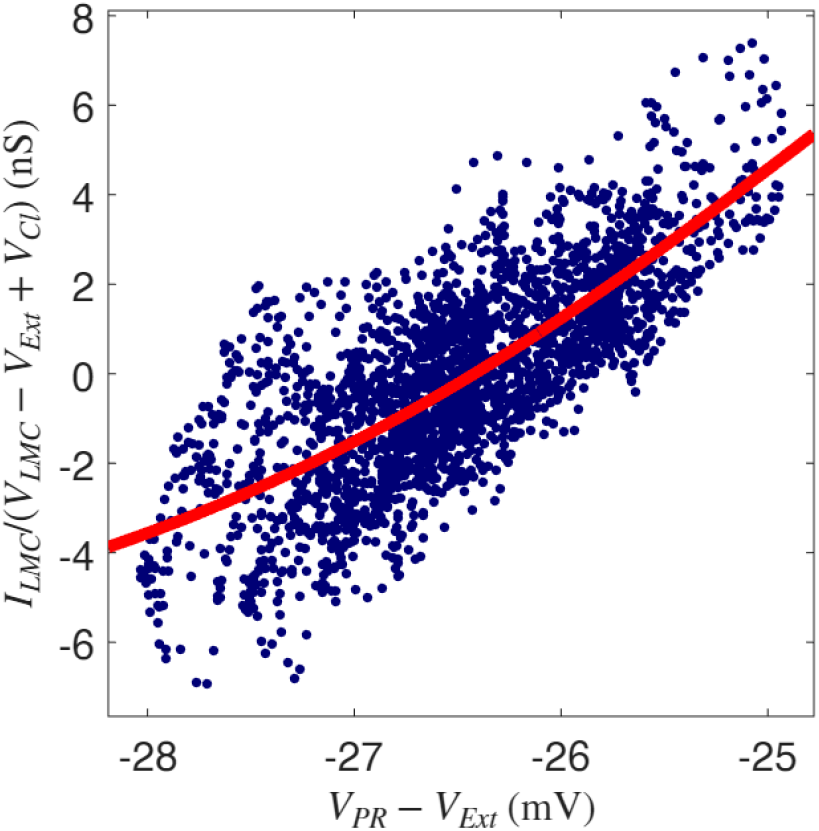
Theoretical conductance of the LMC versus the photoreceptor transmembrane voltage in cartridge 3. Shown in red is the best logistic fit (with a constant shift in conductance *g*_2_, see Equation 9). The slope parameter of the fit is Φ = 1.89mV.

The vesicle release rate is plausibly described by a logistic dependence on the presynaptic transmembrane voltage [16, 28]. Accordingly, we fit the function

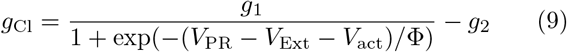

by minimizing the least-squares error, yielding bestfit parameters *g*_1_ = 27.11nS, Φ = 1.89mV, *V*_act_ = *−* 24.53mV and *g*_2_ = 7.31nS. The fitted curve is shown in red. The extra constant *g*_2_ accounts for the fact that *I*_LMC_ only represents AC fluctuations about some mean. The *R*^2^-value of the resulting fit is 0.59. The estimated slope parameter Φ is close to the value Φ = 1.79mV^*−*1^reported by Laughlin et al. (1987) [16] for the logistic relationship between photoreceptor and LMC signal amplitudes across light levels. However, higher values of Φ are obtained in other cartridges, with Φ = 2.70mV and Φ = 2.50mV for cartridges 1 and 2 respectively. In all cases, the data is concentrated near the lower portion of the logistic curve, where the response is effectively exponential.

How does a time-varying photoreceptor voltage enter this feedback framework? Here, we explore the interplay between photoreceptor and extracellular voltages and how these drive vesicle release using a highly simplified toy model. In the idealized scenario of Figure 13, every time the extracellular potential is at or below the photoreceptor potential (minus an activation voltage, which we ignore since it is a constant), a vesicle is released, causing a momentary increase in chloride conductance, which in turn causes the extracellular potential to rapidly increase (due to Cl^*−*^ moving out and into the LMCs) and then relax through impedance to ground. This would happen if the logistic function in Figure 11 and Equation 9 was simplified as a step function. Let us assume that the relaxation is in the linear regime till the next threshold crossing. If the amount *V*_Ext_ rises after the release of a vesicle is some variable amount *a*, the local rate of change of the photoreceptor potential, given the time to the next vesicle release *T* is small (keeping the relaxation dynamic in the linear regime), is given by

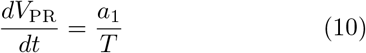

and

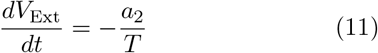

where *a*_1_ and *a*_2_ are as described in the figure. Let us call the difference between the two potentials (i.e., the transmembrane potential) 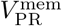. Then,

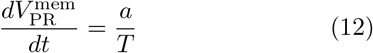

showing, for this simple model, that the rate of vesicle release (and the resulting time-averaged Cl^*−*^ current) is proportional to the local rate of change of the membrane potential.

**FIG. 13.**
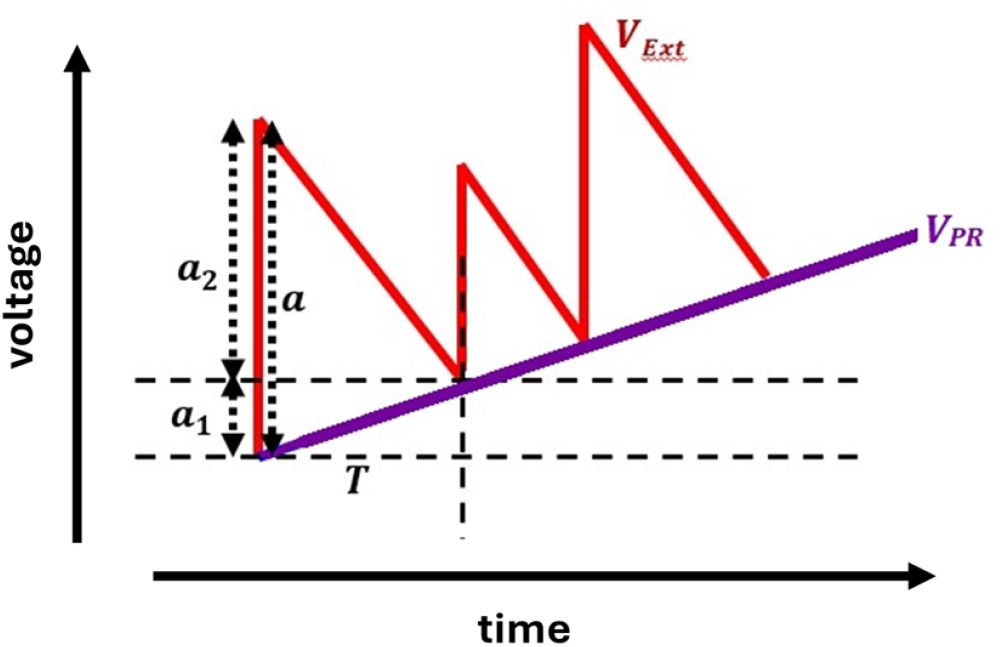
A simple model to show the origin of differentiating behavior in the vesicle rate (and hence the LMC potential). The extracellular potential goes up some variable amount *a* as a result of synaptic activity when the transmembrane potential is 0 (or some activation voltage). This can be broken up into two bits, *a*_1_ and *a*_2_, as shown. Working backwards, this implies *dV*_PR_*/dt* = ⟨ *a*_1_*/T*⟩ and *dV*_Ext_*/dt* = *−* ⟨ *a*_2_*/T*⟩, giving us 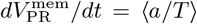 . In other words, the average vesicle release rate tracks the rate of change of the photoreceptor voltage.

While an oversimplified picture, it captures the essential features that show us, at least in part, why we may see differentiating behavior in LMC signals. A rise in photoreceptor voltage shortens the threshold-crossing time for its transmembrane potential (i.e. the time it takes to reach the activation voltage) and hence the time to the next vesicle release, since the photoreceptor voltage more quickly catches up to the increased extracellular potential. The faster the rise, the quicker the arrival of the next event. It thus tells us that when the photoreceptor voltage is changing, the threshold-crossing period of the transmembrane voltage (i.e. the time it takes to reach the activation voltage after the last vesicle release event) depends on the rate of that change. (While not shown here, it is easy to see that conversely, when the photoreceptor voltage *decreases*, the next threshold crossing takes *longer*, and the faster the decrease, the greater the drop in event rate.)

While this interpretation points towards an adaptation based on the electrical properties of the LMCs and the environment of the lamina to improve transmission efficiency, further tests need to be done in order to elucidate what might actually be going on in the synapses. One conundrum already mentioned is that LMC currents are equal and opposite to the extracellular currents during the same light stimulation even though there are *two* LMCs. This discrepancy isn’t explained in any clear way by photoreceptor currents (see Figure 8). Another observation we make is that the known high impedance LMC axon [15, 29] should make it very difficult to draw current up from the ground to create a sufficient feedback voltage in the extracellular space. Both of these issues are solved by thinking of the extracellular impedance as distributed, perhaps because of the tortuosity of the synaptic zone[13, 30], and the recording/current injection site as being separated from the space in the synaptic cleft by a significant impedance. Further analysis of the anatomy and further electrical studies must happen to elucidate to what extent this can really be true.

This work seems to call into question some standard assumptions about quantal synaptic release, particularly the ideas that release events are irregular and statistically independent. However, much of the classic work by Katz and colleagues was conducted in vitro, at relatively quiescent synapses, and relied on invasive pharmacological manipulations [1–3, 31]. Although later studies reported related observations in other systems [3], they also showed that a purely Poissonian description does not fully capture synaptic release dynamics [31–34]. Coordinated vesicle release has been observed, for instance, at specialized ribbon synapses in the vertebrate early visual system, where release events become regularized through subcellular processes such as vesicle–vesicle fusion prior to exocytosis [34]. The functional consequences for synaptic transmission may be similar to what is proposed here. The form of regularization examined in this work (arising from electrical feedback enabled by extracellular electrical isolation in the insect lamina and the sign-inverting nature of the synapse) has not, to our knowledge, been previously explored. It would therefore be interesting to investigate whether analogous electrically mediated regularization mechanisms operate at other synapses.

## IV. CONCLUSION

This study employed a range of analytical tools to investigate visual information transmission in the lamina of *C. vicina*. Analysis of the contrast power transfer and noise power spectral density of voltage responses recorded from photoreceptors and their associated LMCs within the same cartridge, under repeated presentations of a time-varying light contrast signal with a flat spectrum, reveals very high maximum “effective” Poisson rates (Figure 4). While similarly high rates are observed in photoreceptors, the effective rates inferred for LMCs (exceeding *>* 10^5^ per second) are implausibly large, seemingly implying complete turnover of the entire photoreceptor axon membrane several times per second. Notably, the effective Poisson rates measured for the cartridge extracellular potentials in the same experiments are of comparable magnitude to those of the LMCs, suggesting a link between the mechanisms generating the two signals.

Since synaptic activity is fundamentally current driven, we next examined the responses of the recording sites to injected current. The light-induced signals calculated during current injection for photoreceptor, LMC, and extracellular recordings closely match those obtained without current injection, indicating that the responses lie within a linear regime. The resulting current–voltage impulse response functions are consistent with previous RC-fitting results (Tables I, II). The most striking out-come of these experiments is that the light-contrast–to-current impulse response functions are approximately *equal and opposite* (Figures 8, 9). Thus, the same light contrast stimulus induces currents of equal magnitude and opposite sign in the LMC and the extracellular space at the recording site. We interpret these currents as arising from the same synaptic chloride current [24], flowing out of the extracellular space and into the LMC during light-evoked activity, although, as discussed earlier, this interpretation carries important caveats that require further experimental testing. Additional support for a synaptic origin comes from the tight, function-like relationship between the current predicted from the contrastto-current impulse response function and the LMC transmembrane voltage.

High maximum effective Poisson rates can be achieved despite relatively low underlying event rates if those events are regularized through feedback mediated by the extracellular space, which in turn modulates the voltagegated Ca^2+^ channels governing vesicle release. Because each event in this system consumes a finite resource (not only metabolic energy for histamine production but also physical resources such as vesicle availability) the large ratio between the maximum effective Poisson rate and the actual release rate achieved through regularization enhances the efficiency of resource use and accounts for the time-differentiating behavior of the synapse.

The extracellular-feedback–driven regularization mechanism proposed here is primarily qualitative. A more detailed phenomenological model, incorporating the results presented in this work alongside established physiological properties of blowfly photoreceptor–LMC synapses, is developed in Kashef (2025) [28], where this work is originally presented.

## ACKNOWLEDGMENTS

We would like to thank Dr. Sima Setayeshgar for helpful discussions.

## Appendix A Calculating the Effective Poisson Rate

For a flat spectrum signal (eg. a light intensity or photon rate) at time *t*, the contrast function *c*(*t*) tells us by how much the original signal exceeds the average value of the signal as a fraction of that average value (thus, *c*(*t*) is zero-mean). If the signal happens to be a Poisson rate, *λ*(*t*), we can express it as

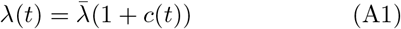

where we always set c(t) to be zero mean, so that 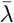 is the average rate. Over multiple repeated trials of applying the same *λ*(*t*) to generate Poisson event sequences, we get an ensemble of event sequences, one of which, when written out as a delta function train, is given by

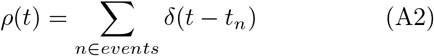

In a measured output signal, the events themselves are not recorded. Rather, a single event leads to a response (a photon absorption voltage bump for a photoreceptor, and a synaptic current pulse-induced voltage impulse). Let *h*(*t*) be the response function. The output signal for a single trial is given by linearly adding the responses to each event in the event sequence for that trial, i.e.

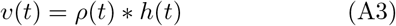

where ‘_*_’ represents convolution. Figure 14 visually represents Equations A1-A3. The ensemble average of *ρ*(*t*) is just *λ*(*t*), so the ensemble average of its fluctuations around the mean is given by

**FIG. 14.**
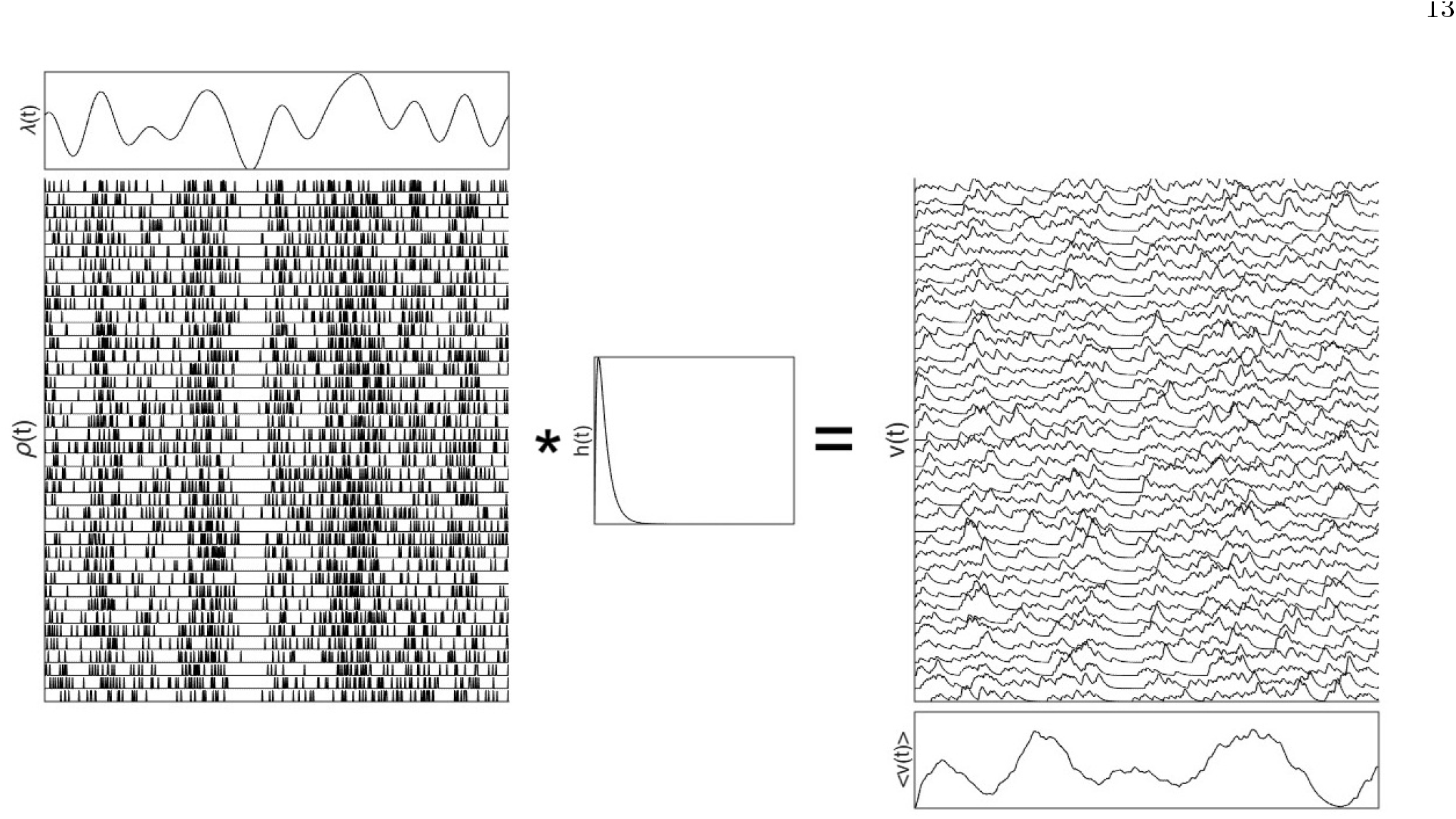
A time varying Poisson rate function *λ*(*t*) gives us, for each trial, an event sequence, which can be represented as a train of delta functions *ρ*(*t*) =Σ _*n∈events*_ *δ*(*t− t*_*n*_) . Convolving the delta function train with an impulse response function *h*(*t*) gives us, excluding added noise, a trial output *v*_*h*_(*t*) = *ρ*(*t*) *\*h*(*t*). Many such event sequences generated over multiple trials from the same *λ*(*t*) can each be used to produce trial outputs in a similar manner, and the ensemble average gives us the output “signal” function *V*_*h*_(*t*) = *⟨v*_*h*_(*t*)*⟩*. The noise trace corresponding to each *v*_*h*_(*t*) (not shown) is then *ε*_*h*_ = *V*_*h*_(*t*) *− v*_*h*_(*t*).

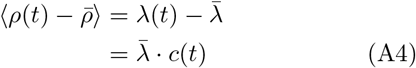

which, combined with Equation A3, gives us

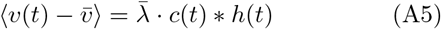

Applying the Fourier transform to Equation A4 gives us 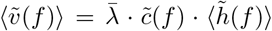 for nonzero frequencies. This leads us to write the *contrast power transfer function* of the output signal as

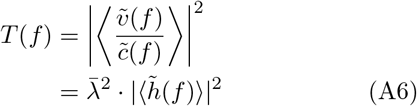

The noise trace for each trial is given by

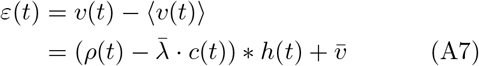

The *noise power spectral density* of the output is thus

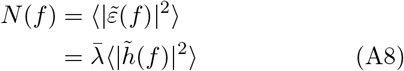

It is the ratio of the contrast transfer function and the noise power spectral density that gives us the effective Poisson rate. Specifically,

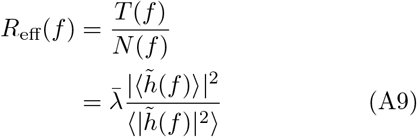

For a noiseless impulse response to an event, 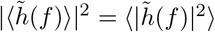, and 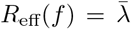, meaning this is exactly the average event rate. When fluctuations are present in the response, per Jensen’s inequality, we always have 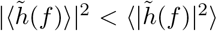, meaning, very generally, 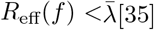 All of this is explained in more detail in de Ruyter and Bialek (2001)[9].

In real life datasets, the calculated *R*_eff_(*f*) isn’t expected to be flat. Jitter across trials has the effect of filtering out high frequency components of the signal (since undesirable timing errors only happen across smaller timescales), but not the noise, which causes the calculated *R*_eff_ to be reduced for higher frequencies. Additionally, of course, one must consider the noise sources other than Poisson noise that affect the calculated noise power spectral density. Let us denote the power spectrum density of the added cumulative noise by *W* (*f*). This would cause the denominator in the first line of Equation A9 to be *N* (*f*) + *W* (*f*) instead of just *N* (*f*) when calculating *R*_eff_(*f*) as the ratio of the measured contrast power transfer and the measured noise power spectral density, thus lowering this calculated *R*_eff_(*f*). For example, 1/f noise, that often appears in precision electronics (see, for example, the photoreceptor and extracellular noise power spectral densities in Figure 4), enhances the noise power spectral density in low frequencies. This lowers the *R*_eff_(*f*) in this part of the spectrum. This is a reason for stating, assuming the underlying process is Poissonian, that the actual rate has to be *at least* the maximum value on the *R*_eff_ plot.

While this method works as shown for estimating the average photon rate during the presentation of a light contrast (in the case of photon flux modulations we know the proportion, c(t), by which the flux is modulated), for LMC signals, this number cannot be directly identified with the supposed vesicle release rate. One has to describe modulation of the release rate by a gain factor *g* (possibly frequency dependent) that converts photoreceptor voltage into vesicle flux (see Laughlin et al. 1987,[16]). If *g* is high, then the synapse encodes relatively reliably at a low mean vesicle rate, but the operating range will be limited by the fact the vesicle rate cannot go below 0. If *g* is low, however, the operating range is large, but the transmission is less reliable. The estimated vesicle rate is the measured *R*_eff_ scaled by a factor of |*g*| ^2^, and release is shut down below contrast values of *−* 1*/g*. Thus if the LMC modulation range is limited to 50%, which is a reasonable estimate as argued in de Ruyter and Bialek (2001) [9], the lower bound on the estimated vesicle release rate is 4 times smaller than *R*_eff_ calculated as in Equation A9.

